# PromoterPredict: sequence-based modelling of *Escherichia coli* σ^70^ promoter strength yields logarithmic dependence between promoter strength and sequence

**DOI:** 10.1101/287607

**Authors:** Ramit Bharanikumar, Keshav Aditya R. Premkumar, Ashok Palaniappan

## Abstract

We present PromoterPredict, a dynamic multiple regression approach to predict the strength of *Escherichia coli* promoters binding the σ^70^ factor of RNA polymerase. σ^70^ promoters are ubiquitously used in recombinant DNA technology, but characterizing their strength is demanding in terms of both time and money. Using a well-characterized set of promoters, we trained a multivariate linear regression model and found that the log of the promoter strength is significantly linearly associated with a weighted sum of the –10 and –35 sequence profile scores. It was found that the two regions contributed almost equally to the promoter strength. PromoterPredict accepts –10 and –35 hexamer sequences and returns the predicted promoter strength. It is capable of dynamic learning from user-supplied data to refine the model construction and yield more confident estimates of promoter strength.

**Availability:** Open source code and a standalone executable with both dynamic model-building and prediction are available (under GNU General Public License 3.0) at https://github.com/PromoterPredict, and require Python 2.7 or greater. PromoterPredict is also available as a web service at https://promoterpredict.com.

## INTRODUCTION

The primary *E. coli* promoter-specificity factor and the one widely used in recombinant DNA technology is the σ^70^ factor. Promoters recognized by σ^70^-containing RNA polymerase are called core promoters and share the following features: two conserved hexamer sequences, separated by a non-specific spacer of ideally 17 nucleotides. The two hexamers are located ~10 bp and ~35 bp upstream of the transcription start site, and are called the – 10 and –35 sequences respectively (Paget and Helmann, 2003; Kadonaga, 2012). Promoters with –10 and –35 sequences matching the consensus motif of the hexamers are typically stronger, meaning they initiate more transcripts per unit time than promoters with less canonical –10 and –35 regions. It is known that the conserved hexamer regions are vital for recognizing and optimizing the interactions between DNA and the RNA polymerase (Hook-Barnard *et al*., 2006; Feklistov and Darst, 2011; Basu *et al.*, 2014).

Theory has yielded a linear relationship between the total promoter score and the natural log of promoter strength (Berg and von Hippel, 1987). Strength of *E. coli* σ^E^ RNA polymerase promoters were studied by Rhodius and Mutalik (2010), who suggested that a study of core (i.e., σ^70^) promoters of housekeeping genes could be complicated by the additional role of transcription activators and limited data on promoter strengths. The complexity of *E. coli* σ^70^ promoter sequences has been treated from an information theoretic standpoint by Shultzaberger *et al*. (2007). Many resources are available to predict the location of promoters in a genomic seqeunce mainly by identifying the −10 and −35 regulatory sequences (for example, de Jong *et al*. (2012)), but there is no (freely) available tool to predict the strength of such sequences. Here we provide a web based platform as well as a standalone tool for the predictive modelling of the strength of σ^70^ core promoters, with the option to dynamically include user data into the predictive model.

## MATERIALS AND METHODS

### Generative model of promoter sequences

A generative model of the −10 and −35 promoter sequences is constructed using two Position Weight Matrices (PWM_−10_ and PWM_−35_) in the following manner. The training set is drawn from the well-characterised Anderson collection of 19 activator-independent promoters maintained at the Registry of standard biological parts (http://parts.igem.org/Promoters/Catalog/Anderson). Nucleotide-wise counts at each position of the hexamer motifs were augmented by a pseudo-count prior to correct for *E. coli* GC content of 50.8% and the resulting frequency matrices were converted into log-odds matrices using Biopython (www.biopython.org).

### Linear modelling of promoter strength

Following Berg and von Hippel (1987), we modelled the relationship between the promoter sequences and the *ln* of the promoter strength using multiple linear regression. Each promoter sequence is scored with respect to the generative models of the −10 and −35 motifs (i.e., the PWM_−10_ and PWM_−35_ matrices) and the two scores obtained formed the feature space of the regression modelling. The regression coefficients to be determined represent the weights of the −10 and −35 regions in the regression analysis. The Anderson promoter library provided promoter strengths normalized in the range 0.00 to 1.00 with respect to the strongest promoter. It was noted that the normalisation step would not affect a linear relationship, altering only the constant of the regression. The normalised strength values were log-transformed to obtain the required response variable values. Since the *ln* function rapidly descends towards – Inf with decreasing promoter strength, we capped the infimum of promoter strength at 0.01 prior to log-transformation. The least-squares cost function was minimized using iterative gradient descent. The model parameters were assessed using t-statistics, and the overall model was assessed using F-statistic and the adjusted multiple coefficient of determination given by:

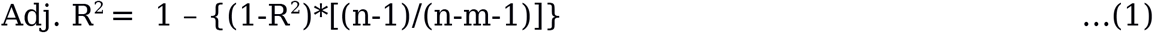

The model was validated using leave-one-out cross-validation (LOOCV).

## RESULTS AND DISCUSSION

**The conservation profile of the −35 and −10 hexamer sequences of the promoters in the Anderson library was visualized using sequence logos and shown in Fig. 1. The site scores of each** promoter sequence were regressed on the ln of the promoter strength. A summary of this process with the training data, log-transformation of the promoter strength and predicted response values is presented in Table 1. The modelling process converged within 10^5^ iterations by tuning the gradient descent to a learning rate (α) of 0.015, and the following model was obtained:

**Table 1.** The promoter activities (strengths) are seen to span the range [0.0, 1.0]. ***\**** indicates promoter strength capped at 0.01. The promoters follow the naming in the Anderson dataset.

| Promoter | PWM<sub>-35</sub> score | PWM<sub>-10</sub> score | Promoter Activity | ln(Promoter Activity) | Predicted ln(promoter activity) |
| --- | --- | --- | --- | --- | --- |
| BBa_J23100 | 8.80192196 | 7.05252226 | 1 | 0 | -1.4669153 |
| BBa_J23101 | 8.50436801 | 8.65655364 | 0.7 | -0.35667494 | -0.25855671 |
| BBa_J23102 | 8.94341939 | 7.79025811 | 0.86 | -0.15082289 | -0.62881141 |
| BBa_J23103 | 5.76111212 | 7.60539274 | 0.01 | -4.60517019 | -4.0308767 |
| BBa_J23104 | 8.94341939 | 8.22340587 | 0.72 | -0.32850407 | -0.22100527 |
| BBa_J23105 | 8.36287058 | 8.22340587 | 0.24 | -1.42711636 | -0.80989265 |
| BBa_J23106 | 8.36287058 | 7.48567002 | 0.47 | -0.75502258 | -1.50446674 |
| BBa_J23107 | 8.36287058 | 8.65655364 | 0.36 | -1.02165125 | -0.40208651 |
| BBa_J23108 | 7.16328158 | 7.91881778 | 0.51 | -0.67334455 | -2.31347961 |
| BBa_J23109 | 8.50436801 | 6.73909721 | 0.04 | -3.21887582 | -2.06383098 |
| BBa_J23110 | 8.36287058 | 7.48567002 | 0.33 | -1.10866262 | -1.50446674 |
| BBa_J23111 | 8.80192196 | 7.48567002 | 0.58 | -0.54472718 | -1.05910916 |
| BBa_J23112 | 5.76111212 | 7.60539274 | 0.00* | -4.60517019 | -4.0308767 |
| BBa_J23113 | 5.76111212 | 7.60539274 | 0.01 | -4.60517019 | -4.0308767 |
| BBa_J23114 | 6.96070112 | 7.48567002 | 0.1 | -2.30258509 | -2.92677594 |
| BBa_J23115 | 7.10219855 | 7.48567002 | 0.15 | -1.89711998 | -2.78324614 |
| BBa_J23116 | 8.94341939 | 7.17224497 | 0.16 | -1.83258146 | -1.21066725 |
| BBa_J23117 | 8.94341939 | 7.17224497 | 0.06 | -2.81341072 | -1.21066725 |
| BBa_J23118 | 8.80192196 | 8.22340587 | 0.56 | -0.5798185 | -0.36453507 |

**Figure 1.**
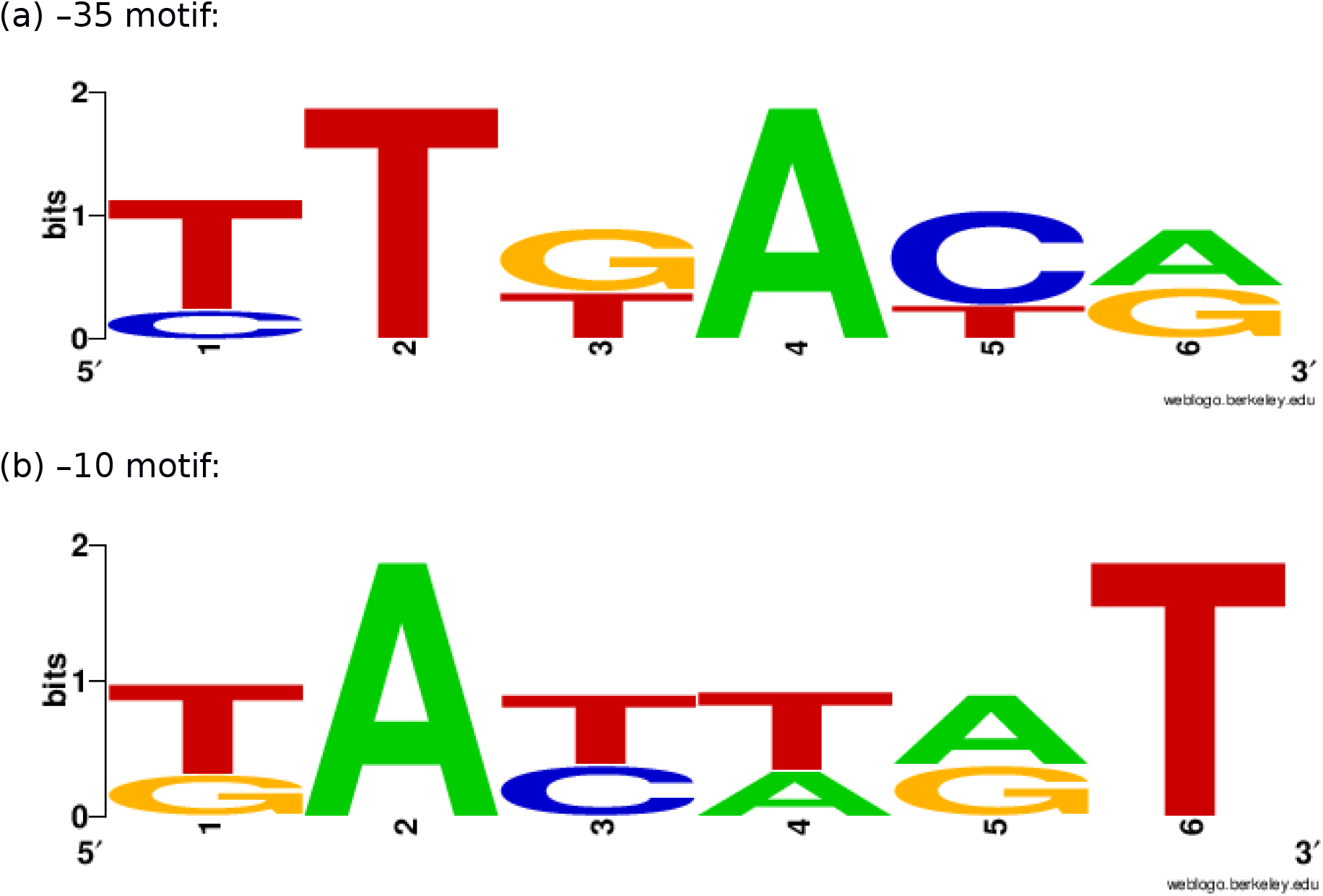
Sequence logos of the −35 and −10 hexamer sequences of the promoters in the Anderson library. Figure was made using WebLogo (Crooks *et al*., 2004).

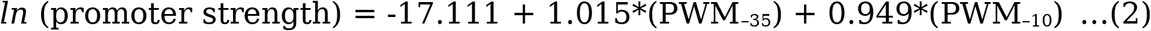

It is observed that the weight coefficients of the two PWM features are almost equal. We derived an independent solution of the multiple regression using `R` (www.cran.org) and obtained a correlation coefficient of 0.998 between the fitted values of the two models. The interval estimates of the coefficients of the regression were computed in `R` using `confint(fit, level=0.95)`, and obtained the following 95% confidence intervals:

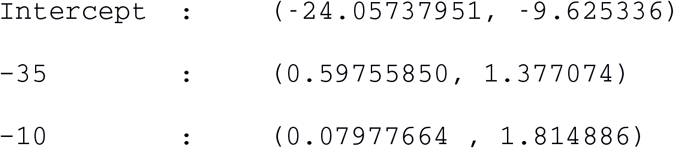

The interval estimates did not include zero, and this implied that the coefficients were significant at the 0.05 level. The p-value of the PWM_−35_ coefficient was < 10^−4^ and that of PWM_−10_ ≈ 0.03. The intercept was significant at a p-value ≈10^−4^. The F-statistic of the overall regression was significant at < 10^−4^ and adj. R^2^ was ≈ 0.65. The plane of best fit corresponding to the above model is visualized in Fig. 2.

**Figure 2.**
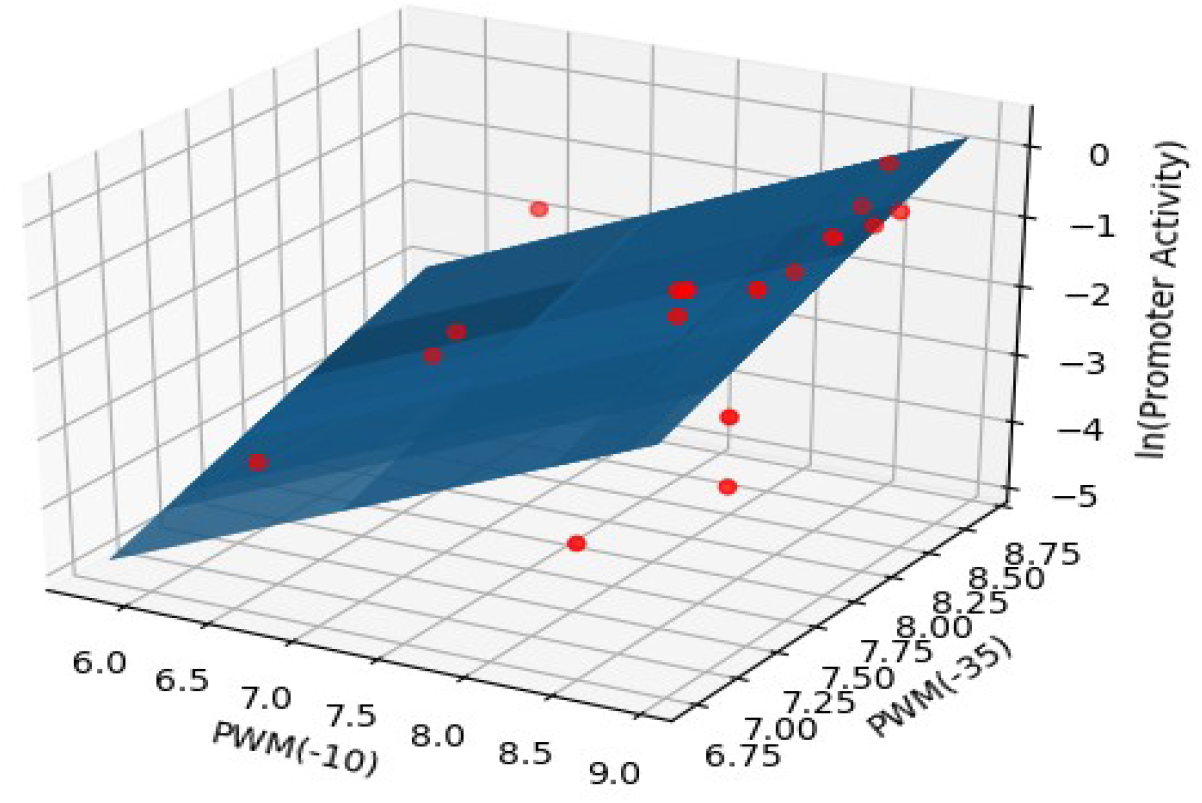
The regression surface (blue) of the estimated model with the training data points(red). X- and y-axes represent PWM scores and the z-axis (vertical) represents the predicted *ln*(promoter strength).

In addition to their independent contributions to promoter strength, we were interested in exploring if any interactions between −35 and −10 sites could contribute to promoter strength. To this end, we tested this possibility in `R` using the following command:

~~~
lm(logStrength ~ PWM35 * PWM10)
~~~

where PWM35 and PWM10 represent the corrresponding site scores. This model resulted in an adj. R^2^ value lesser than that without any interactions. Further, all the four p-values of the regression parameters (intercept, PWM35, PWM10 and interaction) were not significant. The F-statistic was also not significant, thus discounting any interaction between the sites in the present dataset. On this basis, the null hypothesis of absence of any interaction could not be rejected, and we concluded that there is little evidence for interaction between the −35 and −10 sites in determining promoter strength.

Our model assumed that both the predictors carried independent information about the promoter strength, and together they are able to provide sufficient information about the strength. The basis of this assumption was probed to determine if both predictors are necessary to the model. Could one predictor provide sufficient information about the promoter strength in the absence of the other? There are at least three angles to address this question, and all of them were considered to interpret the model better.

(1) Comparing the multiple coefficient of determination with the adjusted multiple coefficient of determination. For the original model:

R^2^ = 0.69

Adj. R^2^ ≈ 0.65

Since there is not much difference between R^2^ and adj. R^2^, we could say that both predictors contribute substantially to the response variable (promoter strength) and account for more than 65% of its variance.

(2) Since the p-values of both predictors are significant, it would be interesting to observe their effect on the response variable in more detail. This was performed using the effects package in `R`:

~~~
library(effects)
fit = lm(logStrength~ PWM35+ PWM10, data)
plot(allEffects(fit))
~~~

The results are shown in Fig. 3. Confidence in the effect of −35 site increases with the −35 score, as evidenced by decreasing uncertainty in logStrength. Such an effect is however not observed for −10 hexamer: the uncertainty widens at both the ends due to edge effects. The effect of the – 35 sequence is also steeper than the effect of the −10 sequence.

**Figure 3.**
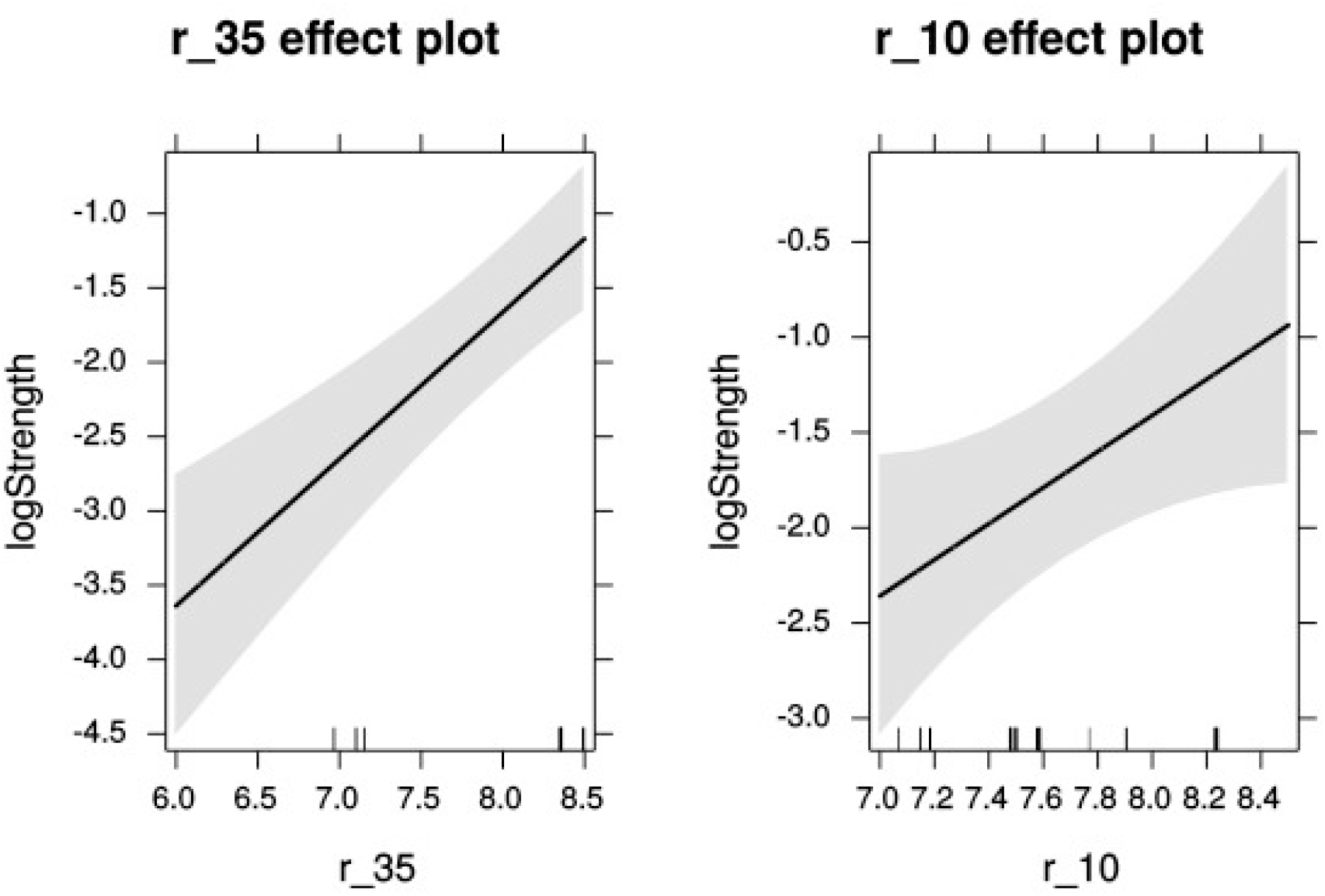
Effects plot of −35 and −10 promoter sites on promoter strength.

(3) Another robust method to address the question is to compute the correlation coefficients between all the variables of interest, including a variable with the combined effects of −35 and −10 sites. This is shown in Fig. 4. Three features were used, namely PWM_−10_ score, PWM_−35_ score, and the combined score. These feature variables were correlated with two response variables, namely promoter strength and its corresponding log transformation. It was first observed that the PWM_−10_ and PWM_−35_ scores were uncorrelated (with a correlation coefficient of just ~0.05). Significantly, the highest correlation between the features and response variable was observed between the combined score and log of the promoter strength (~0.83). This validated our modelling process and was in keeping with similar observations for the strength of σ^E^ promoters (Rhodius and Mutalik, 2010). It was further observed that the combined score showed a relatively moderate correlation with the promoter strength prior to log transformation (about 0.66). This underscored the logarithmic dependence between the promoter strength and sequence, and provided independent validation of Berg and von Hippel’s theoretical model.

**Figure 4.**
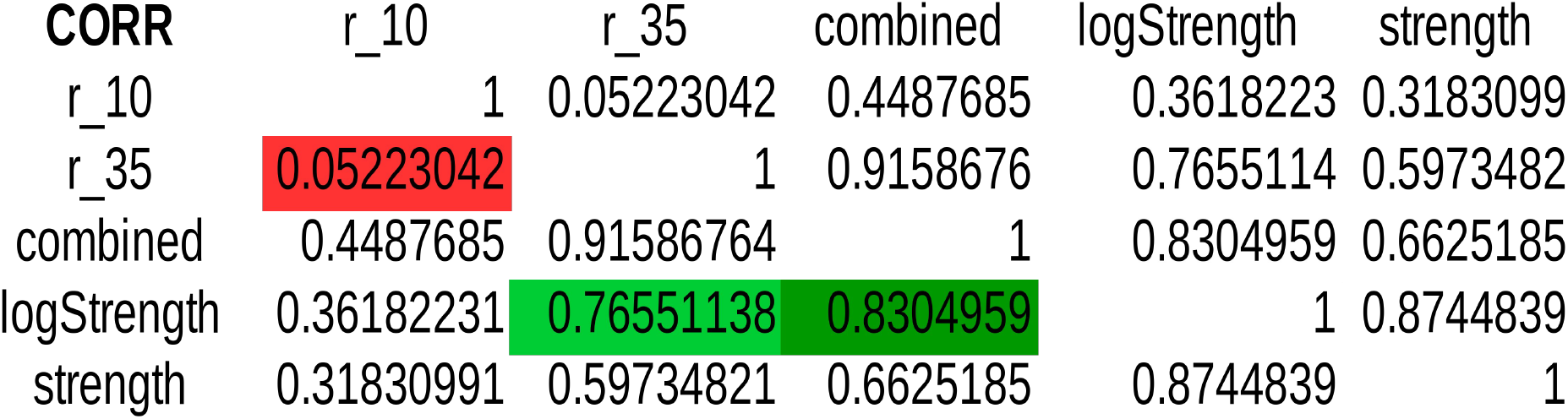
Correlation matrix of features and response variables. Lack of correlation between the predictor variables is highlighted in red. High correlation between features and the response variable is in green.

Finally, the assumptions of linear modelling were investigated with reference to our problem. Model diagnostics of four basic assumptions were plotted (shown in Fig. 5). Specifically:

**Figure 5.**
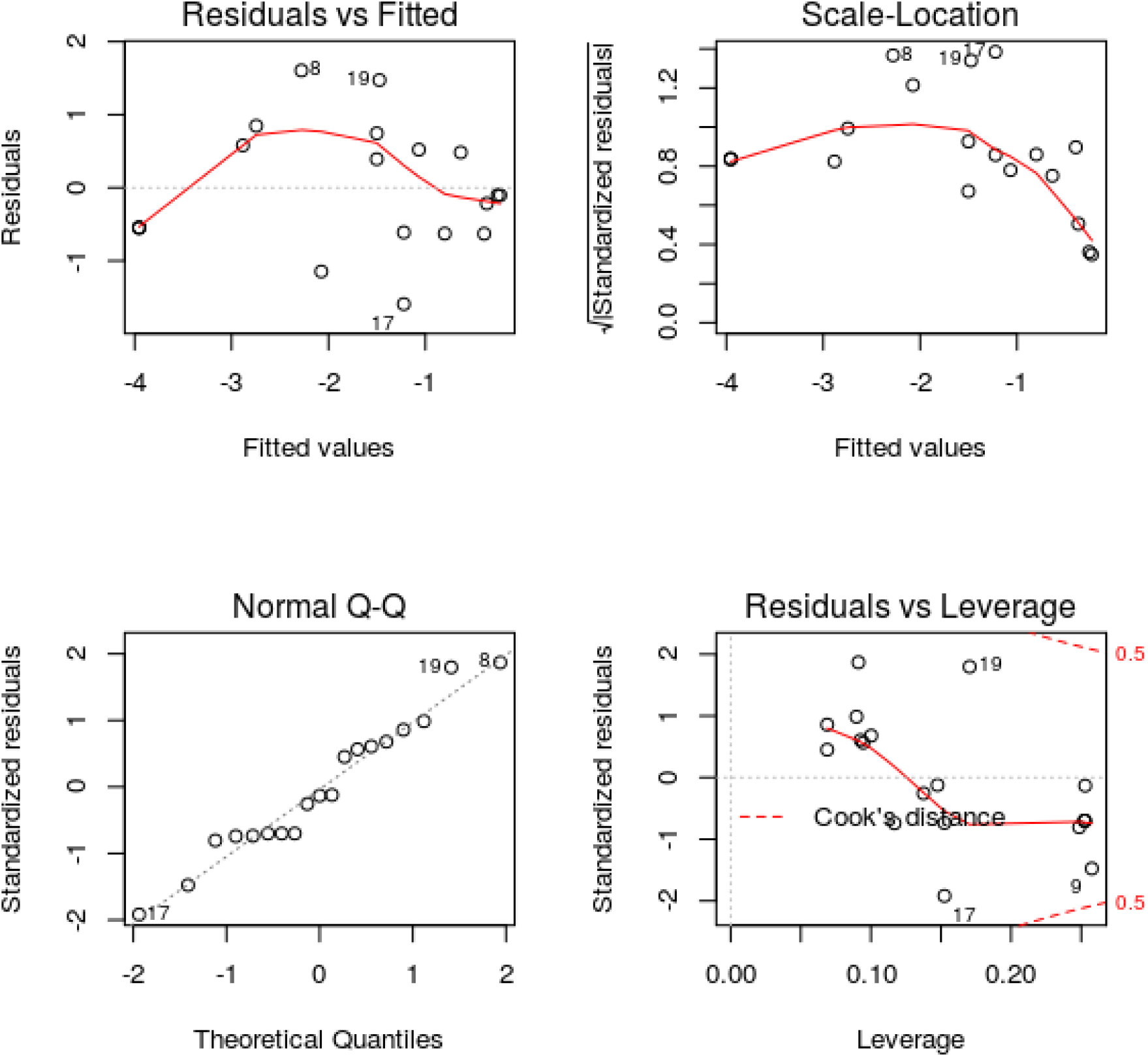
Model diagnostics plots for investigating the assumptions underlying linear modelling. Please see text for discussion.

Plot 1: The residuals were plotted against the fitted values. No trend was visible in the plot, indicating the residuals did not increase with the fitted values and followed a random pattern about zero. This validated the assumption that the errors were independent.

Plot 2: The square root of the relative error (standardized residual) was plotted against the fitted value. No distinct trend was observed, indicating that the standardized residual was not a function of the fitted value. This further validated the assumption that the errors were independent.

Plot 3: To test the assumption that the errors were normally distributed, the standardized residuals were plotted against the theoretical quantiles of a normal distribution. The residual distribution did not significantly deviate from the theoretical quantiles.

Plot 4: Since the least-squares cost function is sensitive to outliers, the number of outliers should be kept to a minimum. This was investigated by plotting the standardized residual against the corresponding instance’s model leverage. This plot showed that there were no significant outliers in the dataset that could exert an undue influence on the regression parameters.

The assumptions of linear modelling were found to be valid, and the model was then cross-validated using a 19-fold LOOCV (similar to jack-knife). Cross-validation yielded a high correlation coefficient of 0.75 (Table 2).

**Table 2.** Cross-validation results. In each trial, a random observation was chosen as a test instance for prediction based on a model built with the rest of the dataset. This process was repeated 19 times, once for each test instance and the cross-validation (CV) residuals were obtained.

| Fold | Observation | Log(strength) | Predicted | CV Pred | CV Residual |
| --- | --- | --- | --- | --- | --- |
| 1 | 3 | -4.600 | -3.995 | -3.771 | -0.729 |
| 2 | 15 | -1.897 | -2.745 | -2.829 | 0.932 |
| 3 | 1 | -0.357 | -0.255 | -0.220 | -0.136 |
| 4 | 10 | -1.109 | -1.501 | -1.530 | 0.422 |
| 5 | 12 | -4.600 | -3.955 | -3.771 | -0.729 |
| 6 | 2 | -0.151 | -0.635 | -0.686 | 0.535 |
| 7 | 4 | -0.329 | -0.228 | -0.210 | -0.118 |
| 8 | 00 | 0.00 | -1.47 | -1.78 | 1.780 |
| 9 | 14 | -2.303 | -2.884 | -2.948 | 0.646 |
| 10 | 5 | -1.427 | -0.800 | -0.717 | 0.710 |
| 11 | 13 | -4.600 | -3.955 | -3.771 | -0.729 |
| 12 | 7 | -1.022 | -0.393 | -0.185 | -0.837 |
| 13 | 17 | -2.813 | -1.222 | -0.936 | -1.877 |
| 14 | 8 | -0.673 | -2.279 | -2.440 | 1.766 |
| 15 | 9 | -3.22 | -2.07 | -1.68 | -1.540 |
| 16 | 11 | -0.545 | -1.067 | -1.120 | 0.576 |
| 17 | 16 | -1.83 | -1.22 | -1.11 | -0.720 |
| 18 | 6 | -0.755 | -1.501 | -1.557 | 0.802 |
| 19 | 18 | -0.580 | -0.366 | -0.332 | -0.248 |

An alternative univariate regression model using only the combined PWM scores found the coefficient to be significant (p-value <10^−4^). However, the weights of the PWMs were slightly different in the model equation (eq. (2)), further the uncertainty in their effects were different. The original multiple linear regression model was retained for the estimation of the promoter strength.

We implemented our model in Python (www.python.org). Since the modelling results are dependent on the dataset, our implementation provides a facility to augment the learning based on user-provided inputs. A web service for the same has been initiated. The web interface is based on Python web module (web.py) and nginx server. The computational layer is based on numpy, Biopython and matplotlib. The user is provided with an option to add any number of promoter instances with −10 and −35 sequences and the corresponding strengths to augment the training data of the supervised model. The goodness of fit of the updated model is re-computed, along with a 3D plot of the regression surface. Based on the trained model, the user could predict the strength of any uncharacterised promoter given its −10 and −35 hexamers.

## CONCLUSION

The following important conclusions were drawn from our study. (1) Sequence-based modelling yielded a logarithmic dependence between promoter strength and sequence. (2) The −10 and −35 sites were equally important in determining promoter strength. (3) The combined sum of the scores (PWM_−35_ + PWM_−10_) emerged as the single most important predictor of the promoter strength. It is straighforward to extend our methodology to the study of promoters of other sigma factors. Our implementation and web service could be useful in characterizing unknown promoters of newly sequenced genomes as well in the selection of promoters for synthetic biology experiments. The dynamic feature of our implementation would enable users with own data to obtain more reliable estimates of promoter strength. The service will be periodically updated based on the availability of new training instances, user input data and/or models for promoters of other sigma factors.

## Acknowledgments

We would like to thank computing facilities at SASTRA Deemed University for support.

